# Chemical degradation of BTK/TEC as a novel approach to inhibit platelet function and thrombosis

**DOI:** 10.1101/2022.05.30.493973

**Authors:** Justin S. Trory, Attila Munkacsi, Kamila M. Śledź, Lucy J. Goudswaard, Kate J. Heesom, Samantha F. Moore, Behnam Nabet, Varinder K. Aggarwal, Ingeborg Hers

## Abstract

The tyrosine kinase BTK plays an important role in platelet function downstream of GPVI and CLEC2 receptors and has been proposed as a novel target to prevent thrombosis in patients that are at increased risk. However, current clinically approved BTK inhibitors have off target effects and are associated with an increased bleeding risk. In this study, we therefore explored whether BTK can be targeted for degradation in human platelets by using recently developed heterobifunctional molecules that employ the proteasomal system to break down BTK. Here we confirm that human platelets are highly susceptible to BTK degraders with the generic tyrosine kinase degrader TL12-186, and the BTK degraders DD-04-15 and DD-03-171 leading to breakdown of BTK and its closely related kinase TEC, an effect that was prevented by proteasomal inhibitors. Tandem Mass Tag proteomic analysis confirmed high selectivity with TL12-186 degrading BTK/TEC, FAK/PYK2 and FER, whereas DD-04-15 and DD-03-171 degraded BTK/TEC only. GPVI-mediated platelet integrin α_IIb_β_3_ activation, P-selectin expression, and phosphatidyl-serine exposure were largely impaired upon BTK/TEC degradation, with PAR-1-mediated responses left intact. This is the first study to demonstrate that chemical protein degraders can be successfully employed in anucleate human platelets to modulate their function.

## Introduction

Studies on mouse models with megakaryocyte/platelet specific deletion of Bruton’s tyrosine kinase (BTK) and/or Tec protein tyrosine kinase (TEC), pharmacological inhibitors, as well as platelets from patients with a functional mutation in BTK, demonstrated an essential role of BTK downstream of ITAM receptors such as GPVI and CLEC2, whereas having little effect on GPCR-mediated platelet function^1,2^. Oral BTK inhibitors furthermore impair thrombus formation triggered by plaques and collagen, suggesting BTK is a promising target as an anti-platelet drug^3,4^. In addition, BTK contributes to FcRIIa-mediated spontaneous platelet aggregation^5^ and responses to bacteria^6^. However, clinically used BTK inhibitors are associated with bleeding, likely to be due to off target effects. In this study we therefore explored whether platelet BTK can be selectively targeted using chemical degraders named Proteolysis Targeting Chimeras (PROTACs); heterobifunctional molecules that contain a E3 ligand, a linker molecule and a ligand for the protein of interest. These degraders mediate ubiquitination and proteasomal degradation of target proteins and multiple groups have confirmed the presence of a functional proteasomal system in human platelets^7^. We here demonstrate that BTK and TEC can be targeted for proteasomal degradation in human platelets, thereby impairing GPVI, but not PAR-1-mediated platelet function. Our findings confirm that chemical degraders can be successfully used in anucleate human platelets, cells that are unable to resynthesise proteins or genetically manipulated, making this is an exciting and promising new research and therapeutic approach to modulate human platelet function and thrombosis.

## Methods

### Platelet Isolation

Blood was obtained from healthy, drug-free volunteers according to local NHS research ethics approval (20/SC/0222) and the Declaration of Helsinki. Blood was drawn into 4% (w/v) trisodium citrate and Acid Citrate Dextrose (ACD) 1/7 (v/v) and centrifuged at 17min/180g. Platelet-rich plasma (PRP) was collected and supplemented with 2μM PGE_1_/0.02U/ml apyrase before pelleting (650g/10 min). Platelets were washed in CGS (13mM trisodium citrate, 22mM glucose, 120mM sodium chloride) with 0.02U/ml apyrase before resuspension in HEPES-Tyrode (HT, 145mM sodium chloride, 3mM potassium chloride, 0.5mM disodium phosphate, 1mM magnesium sulphate, 10 mM HEPES, pH 7.4) supplemented with 5.5mM D-glucose and 0.02U/ml apyrase at a platelet concentration of 4×10^8^ platelets/ml.

### Incubation with BTK degraders

Washed platelets or PRP were incubated with various concentrations of the BTK degraders TL12-186^8^, DD-04-15^8^ and DD-03-171^9^ at 30°C for the indicated period of time. Platelets were subsequently washed in CGS and resuspended in HT as described (see platelet isolation).

### Western blotting and tandem mass tag (TMT) proteomic analysis

Washed platelets were lysed in 4x NuPage sample buffer followed by SDS-PAGE/immunoblotting and probed with appropriate antibodies, as indicated. For TMT proteomic analysis, platelets were lysed in RIPA buffer containing Roche Mini Protease Inhibitor Cocktail and analysed by TMT mass spectrometry using an Orbitrap Fusion Lumos Mass Spectrometer (Thermo Scientific), with data analysis performed using Proteome Discoverer software v2.4 (Thermo Scientific), as reported previously^10^. Normalised protein abundance values were log2 transformed and the within-person fold change (FC) was calculated by subtracting the log2 normalised protein abundance in DMSO from the log_2_ normalised protein abundance in the presence of PROTAC. The mean log2 FC was then calculated for each protein. Paired student’s t-test was used to determine differences in abundance.

### Integrin α_IIb_β_3_ activation and P-selectin expression

Activation of integrin α_IIb_β_3_, α-granule secretion and phosphatidyl serine exposure was assessed in washed platelets by flow cytometry analysis using PAC1-FITC, anti-CD62P-PE and Annexin V-FITC antibodies, respectively, as previously described^11,12^.

## Results and discussion

In this study, we aimed to target BTK for degradation in human platelets using recently developed heterobifunctional molecules that employ the proteasomal system to degrade BTK. Proteomic studies showed that the E3 ligase Cereblon (CRBN), but not Von Hippel-Lindau (VHL), is expressed in human platelets^13^ and CRBN ligands such as thalidomide and pomalidomide were therefore our initial choice of E3 ligase ligands. The Gray lab developed the multi-kinase degrader TL12-186, which contains pomalidomide attached via a linker molecule to the versatile ATP-site directed pharmacophore scaffold TAE684^8,14^. This degrader has previously been shown to potently target BTK for degradation in addition to 27 other proteins depending on the cell line used^8^. The effect of TL12-186 on BTK expression in human platelets was evaluated by Western blotting, demonstrating targeted protein degradation in human platelets (Fig 1A). Incubation of washed platelets with TL12-186 for 4 hours resulted in potent and effective degradation of BTK with maximal degradation reached at 200-500 nM (Fig.1A), confirming that i) chemical degraders are effective in human platelets and ii) BTK is susceptible to protein degradation. Interestingly, higher concentrations of TL12-186 resulted in less effective protein degradation, a ‘hook effect’ phenomenon caused by the three-part system being over-saturated with degrader molecules at higher concentrations^15^. Platelet BTK degradation required over 10 times higher concentrations when incubated in platelet-rich plasma (PRP, Fig.1B), suggesting compound binding to plasma components.

**Figure 1.**
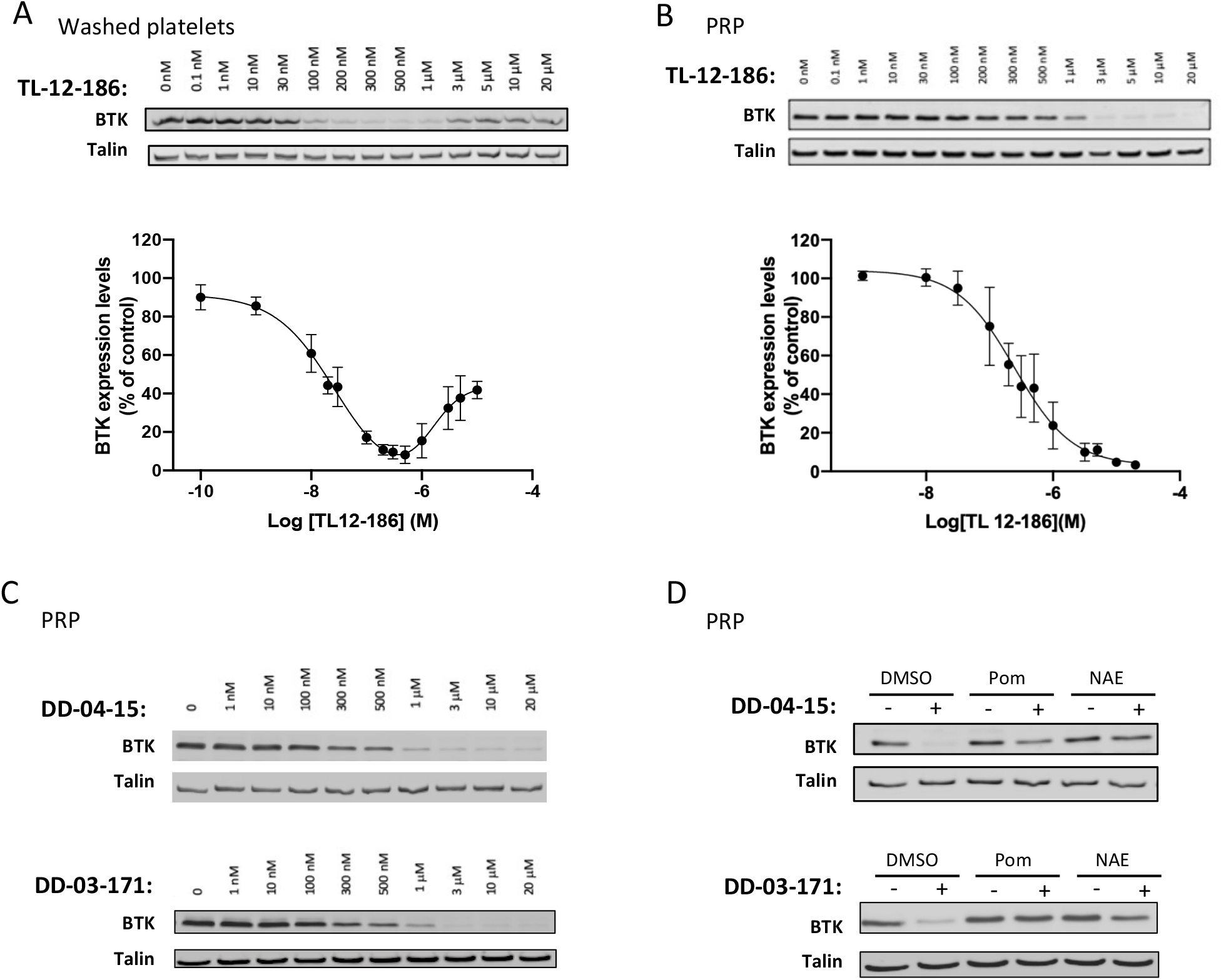
PROTAC-mediated, concentration-dependent degradation of Bruton’s tyrosine kinase (BTK) in human platelets. Washed platelets at 4×10^8^ ml^-1^ **(A)** or platelets in PRP **(B-D)** were incubated with the indicated concentrations of PROTACs 4 or 20 hours/30°C, respectively. In some cases, platelets in PRP were first pre-treated with DMSO, 50 μM Pomalidomide (Pom) or 10μM NEDD8 - activating enzyme (NAE) inhibitor for 4 hours/30°C before incubation with PROTACs for 20 hours/30°C (**D**). Platelets in PRP were subsequently washed in CGS and resuspended in Hepes Tyrodes at 4×10^8^ ml^-1^ (**B-D**). Washed platelets were lysed in 4x NuPage sample buffer containing 0.5M DTT and subjected to SDS-PAGE/immunoblotting with the indicated antibodies (**A-D**). The graphs represent quantification of the results using Odyssey Licor software, expressed as mean fluorescence intensity ± SEM (n=3). Curves (**A,B**) were fitted by non-linear regression using GraphPad Prism 8. A ‘hook effect’ characteristic for the ternary PROTAC degradation system is seen with TL12-186 in washed platelets **(A**), but not with TL12-186, DD-04-015 or DD-03-186 in PRP (**B,C**). Both pomalidomide and NAE prevented BTK degradation **(D)**, confirming E3 ligase/polyubiquitination-mediated degradation. Shown are representative blots (n=3-4; A:3, B:3, C:4, D:3).

As our data demonstrated targeted BTK degradation, we next incubated PRP with the selective BTK degraders; DD-04-15^8^ and DD-03-171^16^. BTK was degraded with similar degradation potency profile to TL12-186 (Fig.1C and SFig.1A). Pomalidomide and NEDD8-activating enzyme (NAE) inhibitor largely abolished PROTAC-mediated BTK proteolysis by DD-04-015 and DD-03-171 (Fig.1D), confirming that BTK is targeted for degradation through the CRBN-mediated ubiquitin proteasomal pathway in human platelets.

To evaluate the specificity of BTK degradation, we implemented a multiplexed quantitative proteomic approach employing TMT. This method allows an unbiased assessment and comparison of the range of targets, as well as the extent of degradation of the proteins of interest by the BTK degraders. TL12-186 selectively degraded BTK/TEC, FAK/PYK2 and FER, demonstrating that these tyrosine kinases can be targeted for PROTAC-mediated degradation in human platelets (Fig.2A,B and Stable1). Interestingly four of these kinases were also degraded by TL-12-186 in the leukemic cell line MOLM-14, but not MOLM-4^8^. The BTK degraders– DD-04-015 and DD-03-171 had remarkable selectivity, with strong degradation of its primary target BTK and TEC, but limited effect on other platelet proteins detected (Fig.2B-D-C, STable 1). Western blotting further confirmed our proteomics results with BTK and TEC being degraded by all compounds, whereas FAK and PYK2 were targeted by TL12-186 only (Fig.2E). The high specificity of the CRBN targeted compounds may be explained by the anucleate nature of human platelets, avoiding changes in protein levels associated with degradation of neo-substrate transcription factors^9^.

**Figure 2.**
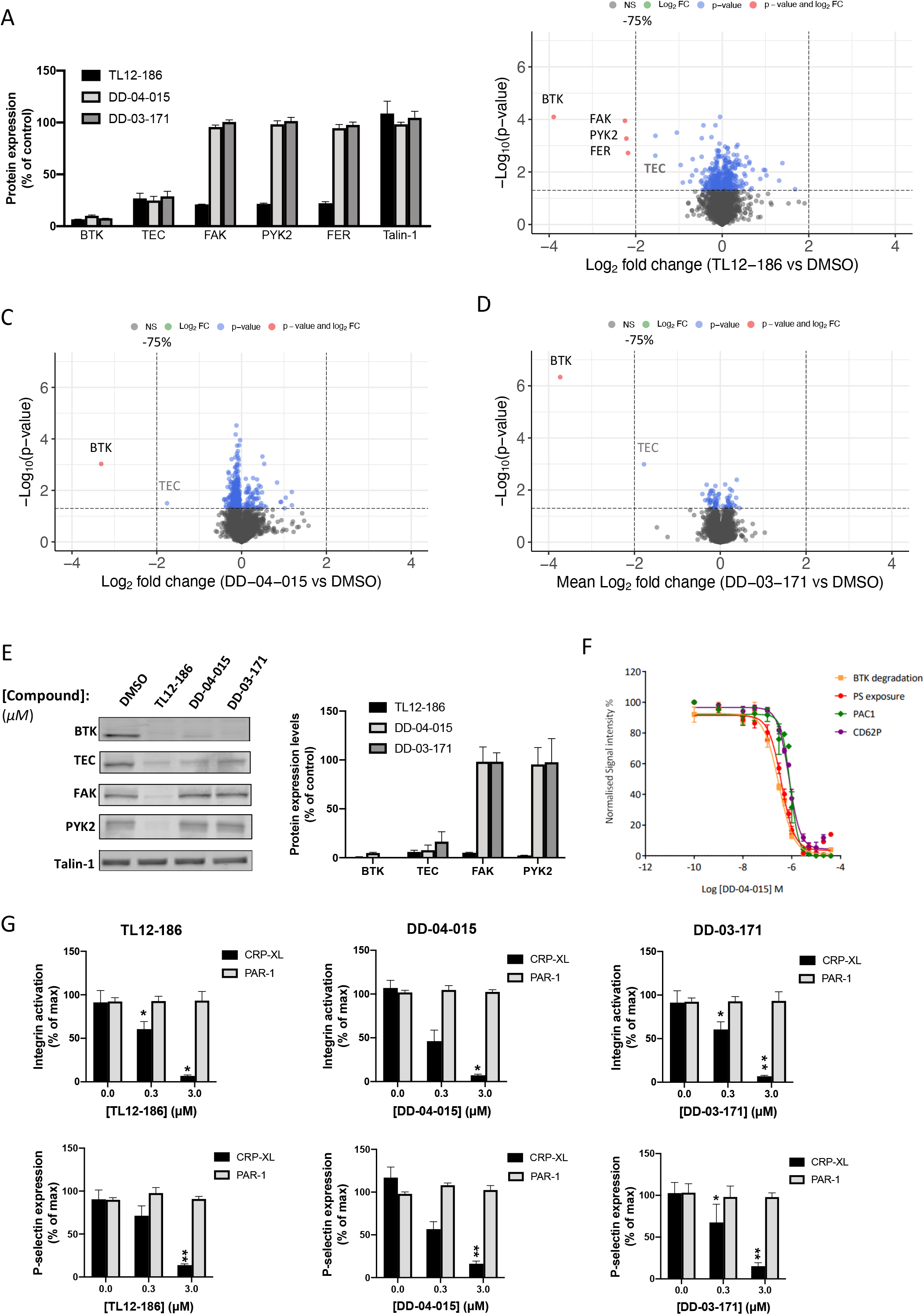
The effect of PROTACs on protein degradation and platelet function. Platelets in PRP were incubated for 20 hours/30°C with either DMSO or with the PROTACs TL12-186, DD-04-015 or DD-03-171 at 3μM (**A-E**) or at the concentration indicated (**F,G**). Platelets were washed in CGS and resuspended in Hepes Tyrodes to 4×10^8^ ml^-1^ for TMT proteomics and western blot analysis (**A-E**, n=3) or to 2×10^7^ ml^-1^ for FACS analysis (**F,G,** n=4). TMT proteomic analysis demonstrates that TL12-186 degrades BTK, TEC, FAK, PYK2 and FER, whereas DD-04-015 or DD-03-171-selectively degrade BTK and TEC **(A)**. Volcano plots of the TMT protein data show the log_2_ fold change in protein expression of PROTAC-treated platelets compared to vehicle (DMSO) treated controls against the -Log_10_ (*p*-value) from a paired student’s t-test. Each dot is representative of one protein. The grey dots represent proteins with a *p*-value >0.05, blue indicates proteins with a *p*-value<0.05 that are <75% degraded. Red indicates proteins with a *p*-value <0.05 that are >75% degraded. TL12-186 degrades BTK, FAK, PYK2 and FER, whereas DD-04-015 or DD-03-171 degrade BTK to >75%, p<0.05. Volcano plots were made using the “EnhancedVolcano” R package. **(B-D)**. TEC is indicated in grey as it did not pass the analytical software internal threshold for TMT quantification (**B-D**). Western blot analysis confirmed the TMT proteomics data. The graphs represent quantification of the results using Odyssey Licor software, expressed as mean fluorescence intensity ± SEM (n=3) **(E)**. DD-04-015-treated washed platelets were stimulated with 2μg ml^-1^ CRP-XL for 10 minutes in the presence of PAC1-FITC/CD62P-PE to assess integrin α_IIb_β_3_ activation and P-selectin expression, respectively. Alternatively, platelets were stimulated by a combination of CRP and thrombin in the presence of Annexin-V-488 fluorescent dye to measure levels of PS exposure. Results are expressed as mean % of maximal normalised signal intensity ± SEM, n=4 (**F**). PROTAC-treated platelets were stimulated with 2μg ml^-1^ CRP-XL or 10 μM TRAP-6 for 10 minutes in the presence of PAC1-FITC/CD62P-PE to assess integrin α_IIb_β_3_ activation and P-selectin expression, respectively. Results are expressed as mean % of maximal normalised signal intensity ± SEM, n=4 (**G**). PROTAC-mediated protein degradation concentration dependently reduced CRP-mediated, but not PAR-1-mediated, integrin α_IIb_β_3_ activation and P-selectin expression. Statistical analysis was performed using a repeated measures one-way ANOVA, *p≤0.05 **p≤0.01 **(F,G)**.

To determine the role of BTK/TEC in human platelets, we evaluated CRP-mediated integrin α_IIb_β_3_ activation, P-selectin expression (a marker of α-granule secretion), and phosphatidyl serine (PS) exposure and compared this to the extent of BTK proteolysis by DD-04-015. Figure 2F demonstrates that degradation of BTK by DD-04-015 compound led to a concentration-dependent inhibition of CRP-mediated integrin α_IIb_β_3_ activation, α-granule secretion, and PS exposure. Interestingly, PAR-1-mediated platelet function was unaffected (Fig.2G), confirming that the BTK degraders specifically target the GPVI pathway. A similar difference was observed in platelets from patients with a functional mutation in BTK^1^. Together, our data shows highly selective degradation of platelet BTK and TEC by protein degraders. We thus demonstrate for the first time the enormous potential of chemical degraders in modulating platelet function and preventing thrombosis. With the PROTAC field growing exponentially and the inability of platelets to resynthesise proteins, PROTAC-mediated protein degradation is an exciting and promising therapeutic approach to modulate platelet function and thrombosis.

## Supporting information

STable1

## Acknowledgements

We thank the healthy blood donors within the University of Bristol for their generous donations and Nathanael S. Gray (Chemical and Systems Biology, Chem-H, Stanford Cancer Institute, Stanford Medicine, Stanford University, Stanford, CA, USA) for providing DD-04-15 PROTAC. This work was supported by NC3Rs, Bristol Alumni and the British Heart Foundation (grant RG/15/16/31758, FS/16/27/32213, PG/16/3/31833, PG/16/21/32083, FS/17/60/33474, SP/F/21/150023, AA/18/7/34219). The graphical abstract was created using BioRender.

## Authorship Contributions

Conceptualization, I.H., S.F.M., and V.K.A; Data curation, I.H. and K.M.H.; Formal analysis, J.S.T., A.M. K.M.S., L.J.G. Funding acquisition, I.H. and V.K.A.; Investigation, J.S.T., A.M. K.M.S. and I.H.; Methodology, J.S.T., A.M., K.M.S., S.F.M., B.N.; Supervision, I.H.; Writing—original draft, I.H.; Writing—review and editing, J.S.T., K.M.S, L.J.G., B.N., V.K.A.

